# A Systems Framework for Quantifying Programmability and Persistence Across Mammalian Cell Types

**DOI:** 10.64898/2026.03.27.714669

**Authors:** Viren Chauhan, Mark Chen, Anusa Thangavelu Sridharan, Lurong Pan

## Abstract

Cellular therapies, toxicity screening, and regenerative medicine depend on selecting mammalian cell types with optimal lifespan, persistence post-transplant, immunogenicity, and chemical resilience. This review synthesizes data from over 50 immune, parenchymal, stem, and emerging engineered cell populations—including gamma-delta T cells, iNKT cells, CAR-macrophages, and hypoimmune iPSC derivatives—drawing from in vivo lifespan studies (including ^1^◼C birth-dating and deuterium labeling), engraftment dynamics, immune rejection risk, and stress sensitivity profiles. We introduce a Programmability & Persistence Score (PPS; 0–20) that integrates these features into a unified metric, complemented by Pareto frontier analysis to visualize multi-objective trade-offs. High-PPS cell types (e.g., HLA-matched HSCs, hypoimmune iPSCs, chondrocytes) are suited for long-term regenerative applications, while low-PPS sentinels (e.g., neutrophils, enterocytes) serve acute assays. We discuss mathematical extensions including multi-criteria decision analysis, fuzzy membership functions, and Bayesian frameworks that address limitations of linear additive scoring. Together, these integrated profiles support cell selection for gene editing, organ-on-chip systems, in vivo cell programming, and immunotherapy, bridging cell biology with translational engineering.

## 1. Introduction

The expanding landscape of biomedical applications—including regenerative grafting, immunotherapy, organ-on-chip design, and synthetic biology—relies critically on the rational selection of mammalian cell types^1^. Yet, decision-making remains ad hoc due to the absence of a quantitative, comparative framework^2^. Recent advances in hypoimmune engineering^228^, in vivo cell programming^229^, and AI virtual cell models^173^ have dramatically expanded the repertoire of candidate cell types, making systematic comparison more essential than ever.

To address this gap, we develop a mathematically grounded systems framework to evaluate cell types using the Programmability & Persistence Score (PPS). The PPS integrates four biologically relevant traits:

- **Intrinsic stability** – determining long-term survival^5^;
- **Post-transplant persistence** – capturing engraftment and functionality^6^;
- **Immunogenicity** – inversely scored to reflect immune compatibility^7^;
- **Chemical resilience** – indicating environmental durability and stress tolerance^7^.

We further introduce Pareto frontier analysis to visualize multi-objective trade-offs that linear scoring may obscure, and discuss extensions through multi-criteria decision analysis (MCDA), fuzzy logic, and Bayesian hierarchical modeling. Through application to over 50 cell types plus emerging engineered populations, we demonstrate the framework’s flexibility and potential for standardizing cell-type evaluation across domains^10–12^.

## 2. Profiles of Immune and Parenchymal Cell Types

### 2.1 Innate Immune Cells

Innate immune cells serve as frontline responders with variable transplant potential^31–33^. Neutrophil lifespan remains debated: conventional labeling yields a circulating half-life of ∼19 hours, while in vivo deuterium labeling suggests 5.4 days including the tissue phase^34^. Koenderman et al. (2022) proposed a “conveyor belt model” where blood neutrophils exchange with a large tissue-resident pool^230^. PPS scoring reflects both estimates with appropriate caveats. Monocytes persist 1–3 days in vivo with high immunogenicity (MHC-II inducible)^38,39^.

Tissue-resident macrophage lifespans vary considerably. Microglia turnover is now precisely quantified through ^1^◼C dating at ∼28% per year (average cell age 4.2 years), with some cells persisting >20 years^231^. Kupffer cells self-renew for months–years with moderate immunogenicity aided by liver tolerogenicity^40–42^.

### 2.2 Adaptive Immune Cells

Adaptive cells offer memory and specificity but pose immunogenicity challenges in allogeneic settings^46–48^. Naïve CD4^+^/CD8^+^ T cells last 2–6 years with very high immunogenicity^49,50^. Central/effector-memory T cells endure months–years with low–moderate sensitivity^53,54^. Regulatory T cells persist years and prolong grafts^55,56^. Long-lived plasma cells may persist for decades to a lifetime^232^.

### 2.3 Emerging Therapeutic Immune Populations

#### Gamma-delta(γδ) T cells

possess MHC-independent antigen recognition, eliminating GvHD risk and enabling off-the-shelf allogeneic products. Adicet Bio’s anti-CD20 CAR γδ T cell received FDA Fast Track designation for mantle cell lymphoma^233^. Their persistence profile differs substantially from αβ T cells, warranting separate PPS scoring.

#### Invariant NKT (iNKT) cells

recognize lipid antigens via CD1d. Cellistic/BrightPath Bio initiated a Phase I trial of iPSC-derived BCMA-targeting CAR-NKT cells in December 2024^234^. **MAIT cells** show unique tumor microenvironment remodeling capability. **CAR-macrophages** (Carisma Therapeutics, first-in-human) offer TME remodeling through enhanced phagocytosis but face persistence limitations inherent to myeloid lineages^235^. **Engineered Tregs** represent a distinct immunosuppressive modality for transplant tolerance. **iPSC-derived NK cells** (Century Therapeutics’ CNTY-101: 83% ORR at high dose) combine off-the-shelf manufacturing with potent cytotoxicity^236^.

### 2.4 Representative Tissue and Parenchymal Cells

For hepatocytes, the landmark ^1^◼C birth-dating study by Heinke et al. (*Cell Systems*, 2022) revealed a critical distinction: diploid hepatocytes have an average age of less than 3 years with >7-fold higher annual birth rates than polyploid hepatocytes^237^. This heterogeneity has direct implications for PPS scoring and is reflected as a range rather than a single value. Cardiomyocyte renewal remains at ∼1% annually at age 20, declining to ∼0.3% at age 75^67,68^. Chondrocytes endure decades with moderate immunogenicity due to their avascular niche^89,90^.

## 3. Stem Cell Profiles: Chemical Sensitivity and Half-Life Rankings

Stem cells vary in potency and fragility. Rankings reflect functional half-life in minimalist 2D culture^98–100^:

1. **HSCs**: <24h (>50% attrition overnight). UM171 (dorocubicel), evaluated in >120 patients, achieves median neutrophil engraftment of 9.5 days versus 22 days for unmanipulated cord blood; received FDA RMAT and EMA PRIME designations^238^. Combination with SR1 sustains HSC culture to day 45^239^.
2. **NSCs**: 3–5 days; sensitive to ROS, glutamate, ER-stress^104–106^.
3. **Satellite stem cells**: ∼6 days (Pax7 loss); extend with soft matrices^107–109^.
4. **MSCs**: 1–2 weeks; sensitive to telomerase limits. Ryoncil (remestemcel-L), the first FDA-approved MSC therapy (December 2024), confirmed paracrine “hit and run” mechanism^240^.
5. **ESCs**: Indefinite in colonies but 40–70% dissociation apoptosis without ROCK inhibitor. CEPT cocktail (Chroman 1 + Emricasan + polyamine + Trans-ISRIB) gaining adoption^241^.
6. **iPSCs**: Indefinite; slightly better single-cell survival than ESC. 115–116 clinical trials testing 83 hPSC products with >1,200 patients dosed as of January 2025^242^.

## 4. PPS Model Construction and Mathematical Extensions

### 4.1 Base Weighted Linear Model

The base PPS defines a weight vector **w** = (w_stab_, w_persist_, w_imm_, w_chem_). The weighted PPS for a specific application is:

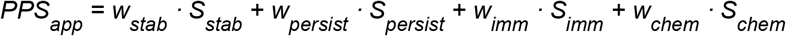

where each S_factor_ is the sub-score (0–5). Weights are normalized (∑w_i_ = 4) so PPS_app_ remains on a 0–20 scale. Cell therapy may use **w** = (1, 2, 2, 1); toxicology PPS may use **w** = (1, 1, 1, 2)^227^.

### 4.2 Limitations of Linear Additive Scoring

The MCDA literature identifies several well-characterized limitations of linear weighted scoring that apply to the PPS methodology^243^. **Compensatory bias**: low scores on one criterion compensate for high scores on another, which may be biologically inappropriate (e.g., high immunogenicity should not be “compensated” by high stability for transplant applications). **Preferential independence**: linear additive models assume criteria are mutually independent, which biological parameters rarely are.

#### Weight sensitivity

small changes in sub-score weights can dramatically alter cell type rankings.

#### Nonlinearity

biological systems exhibit fundamentally nonlinear dose-response relationships.

### 4.3 Pareto Frontier Analysis

We complement the linear PPS with Pareto frontier analysis (Figure 5). Shoval et al. (*Science*, 2012) demonstrated that organisms performing multiple tasks occupy low-dimensional polyhedra in trait space, with vertices as “archetypes” optimal at single tasks. Cell types may represent Pareto-optimal trade-offs rather than being rankable on a single linear scale—a neuron optimizes for lifespan at the cost of renewability, while an enterocyte optimizes the inverse. Mapping cell types onto the programmability-persistence Pareto frontier reveals that HIP-iPSCs, HSCs, and neurons define the frontier boundary, while most immune cells occupy the interior, indicating room for engineering improvement^244^.

**Figure 1.**
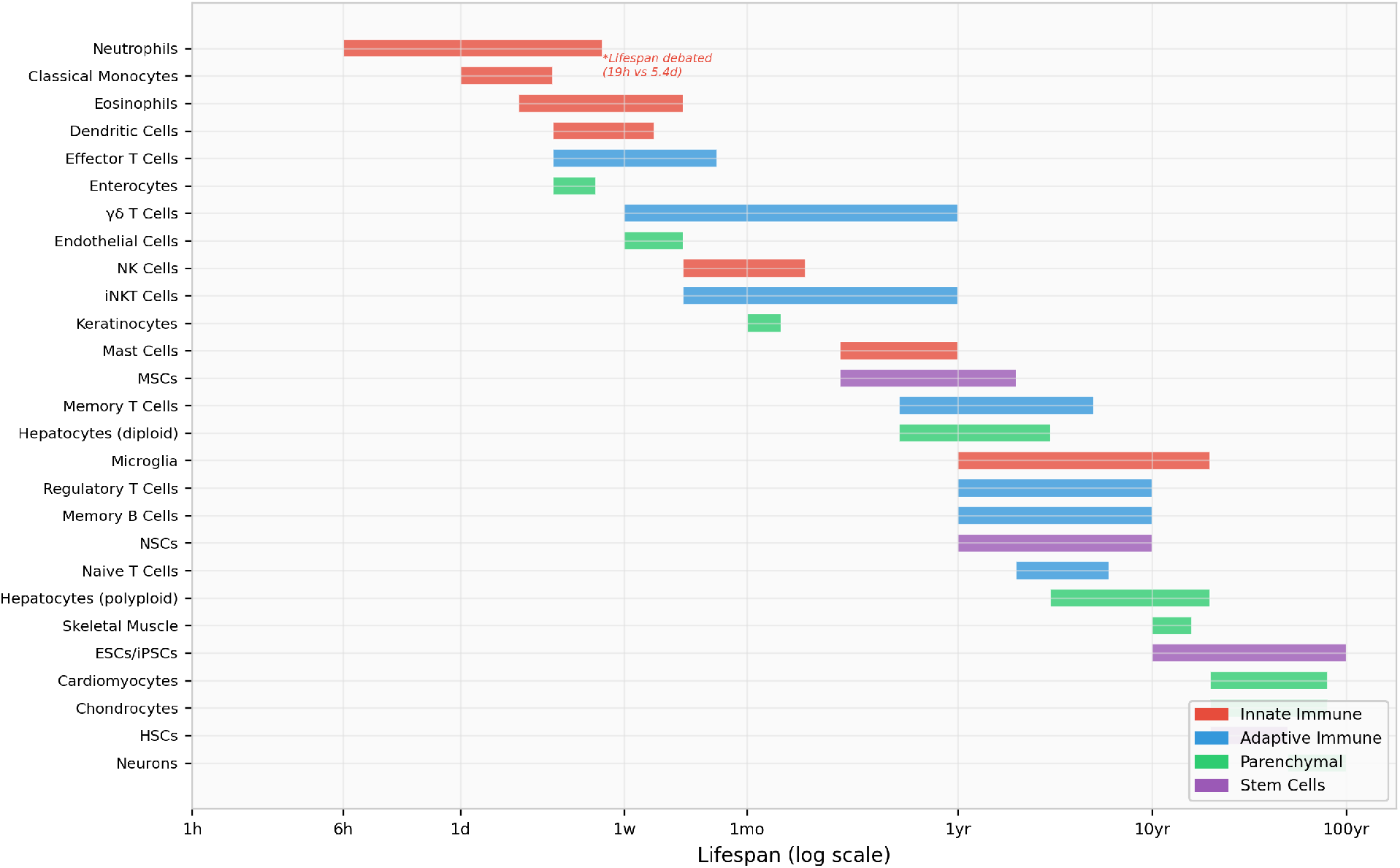
Cell Lifespan Timeline Across Mammalian Tissues (Updated 2025). Horizontal bars represent reported in vivo lifespans on a logarithmic scale. Notable updates include the neutrophil lifespan controversy (19h vs. 5.4d), hepatocyte diploid/polyploid distinction (^1^◼C data), microglia average age of 4.2 years, and inclusion of γδ T cells and iNKT cells.

**Figure 2.**
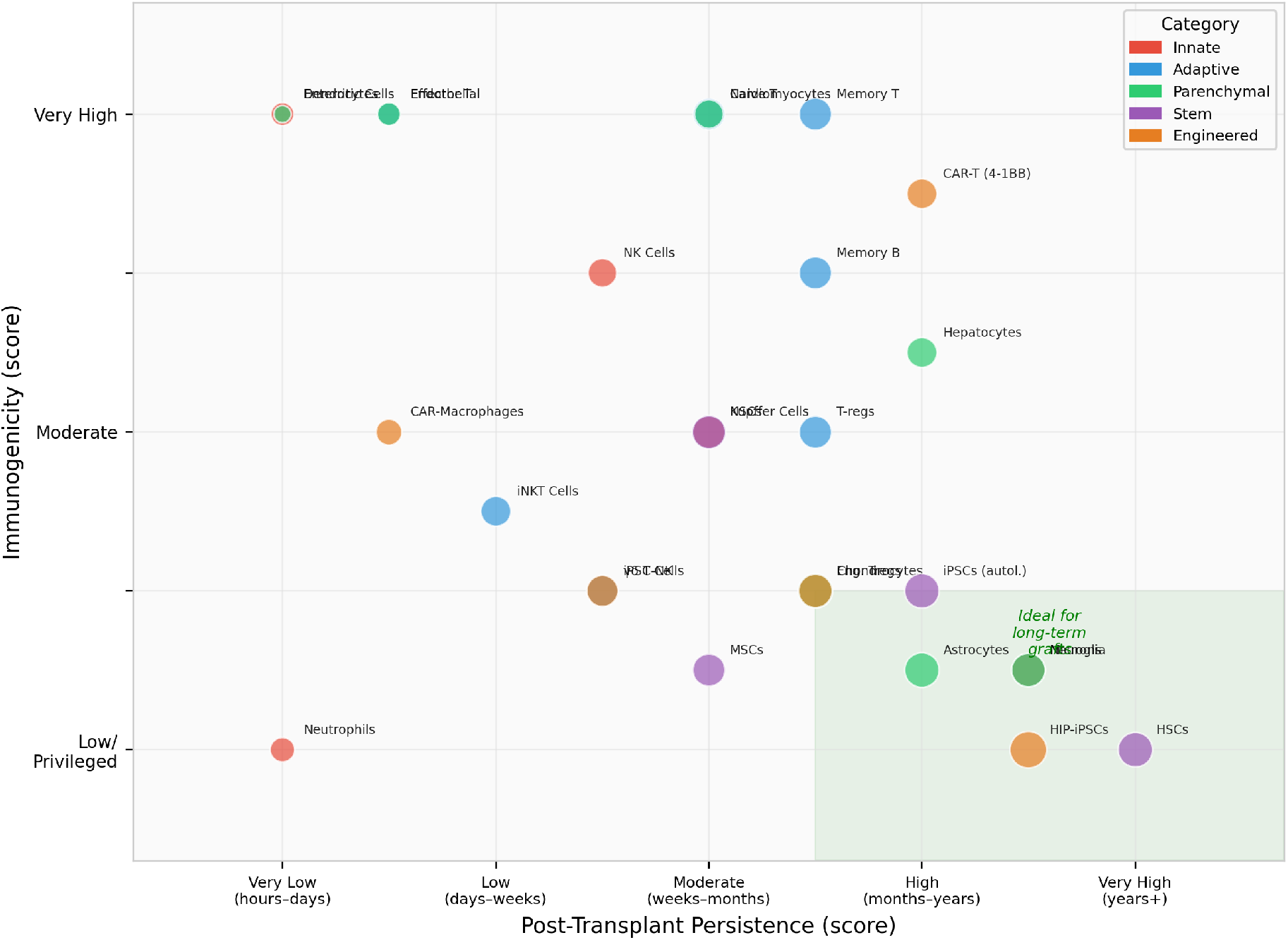
Immunogenicity vs. Persistence (Updated with Engineered Cell Types). Bubble size reflects PPS. Orange markers denote engineered populations including HIP-iPSCs, CAR-T (4-1BB), iPSC-NK, engineered Tregs, and CAR-macrophages. The green-shaded quadrant indicates the ideal zone for long-term grafts (low immunogenicity, high persistence). HIP-iPSCs achieve the highest combined score.

**Figure 3.**
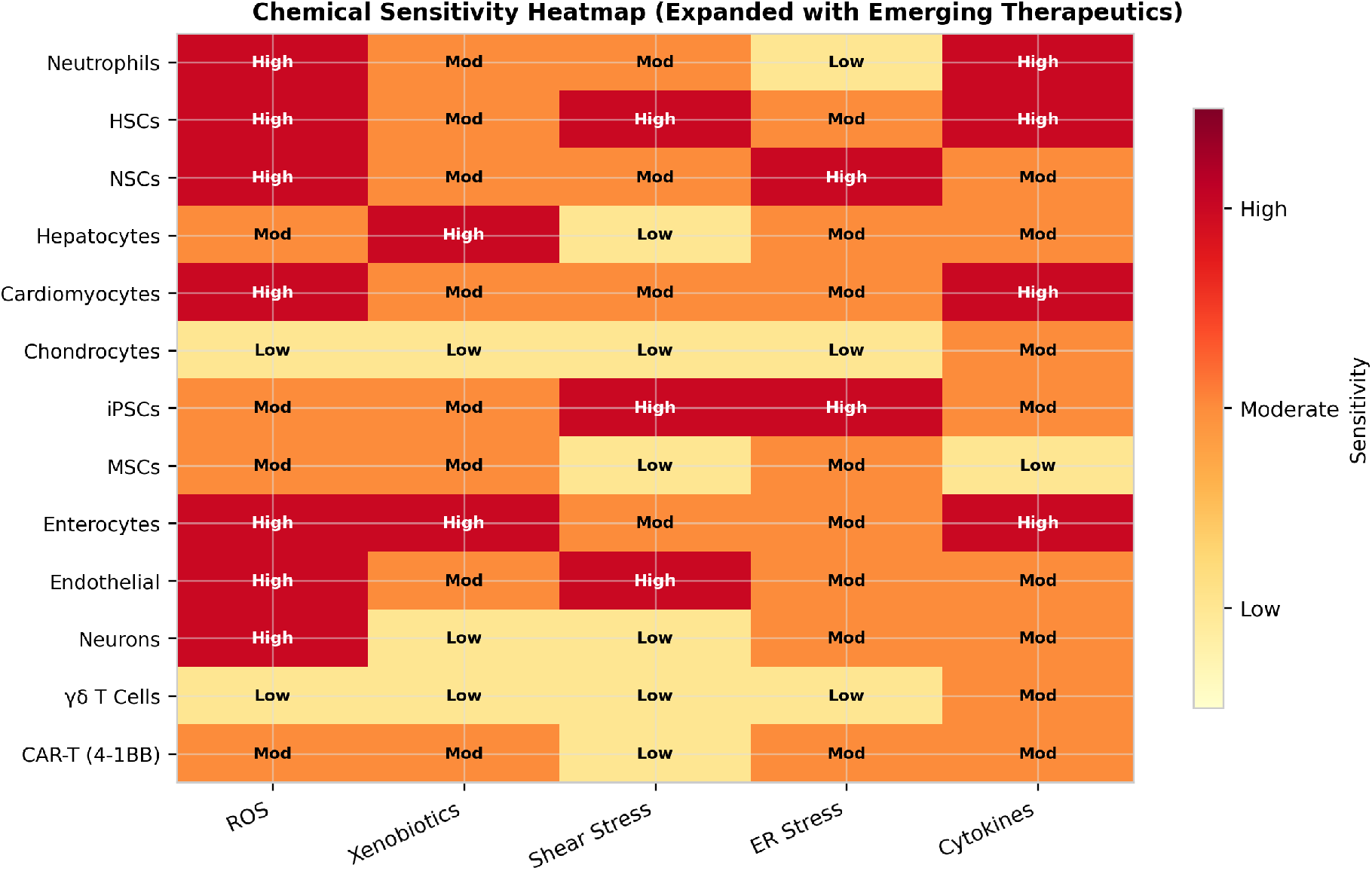
Chemical Sensitivity Heatmap (Expanded). Now includes γδ T cells and CAR-T (4-1BB) products. γδ T cells show broad resilience similar to chondrocytes, supporting their potential as robust off-the-shelf therapeutics.

**Figure 4.**
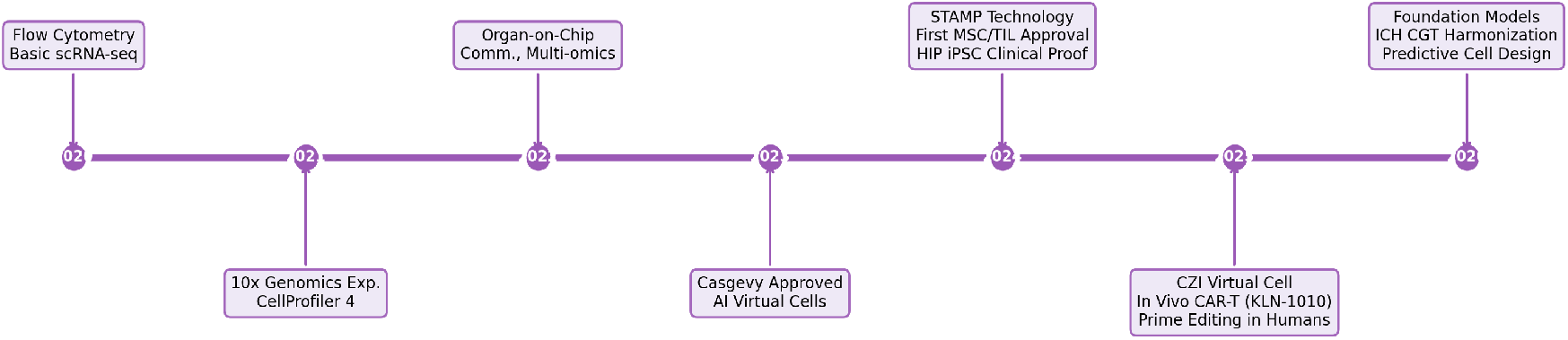
Technology Evolution in Cell Profiling (2020–2026). Updated milestones include Casgevy approval (Dec 2023), first MSC/TIL FDA approvals and HIP iPSC clinical proof (2024), CZI Virtual Cell platform launch, in vivo CAR-T (KLN-1010) first-in-human data, and first prime editing in humans (2025).

**Figure 5.**
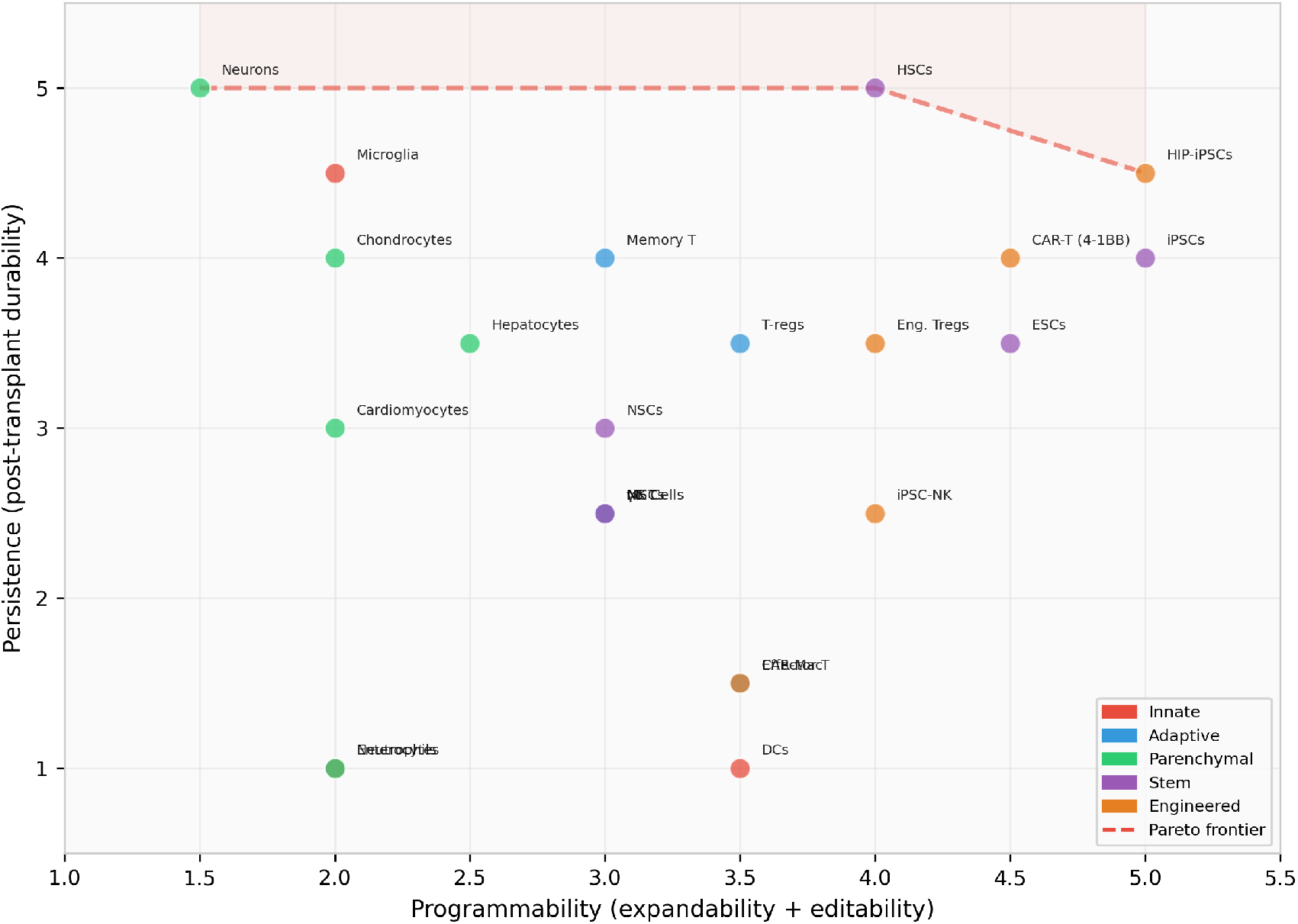
Pareto Frontier Analysis: Programmability vs. Persistence Trade-offs. Cell types are plotted by programmability (expandability + editability) against persistence (post-transplant durability). The dashed red line indicates the Pareto frontier—cell types on or near this boundary represent optimal trade-offs. HIP-iPSCs, HSCs, and neurons define the frontier. Engineered populations (orange) generally shift toward higher programmability relative to their natural counterparts.

### 4.4 Alternative Mathematical Frameworks

**Fuzzy logic** handles inherent biological variability more naturally than crisp scores. Fuzzy membership functions can represent measurement uncertainty (e.g., the neutrophil lifespan controversy) rather than forcing point estimates^245^. **Bayesian hierarchical models** can integrate heterogeneous data sources (clinical trials, preclinical, in vitro) with appropriate uncertainty quantification. **TOPSIS** (Technique for Order of Preference by Similarity to Ideal Solution) ranks cell types based on distance from an ideal programmability-persistence profile, naturally handling multi-criteria structure without assuming linear additivity^243^. We recommend these as complementary analyses in future implementations of the PPS framework.

## 5. Programmability & Persistence Score: Integrated Ranking

The PPS (0–20) sums sub-scores for in vivo stability (decades/lifelong = 5; months–years = 3; ≤days = 1), post-transplant persistence (years/indefinite = 5; weeks–months = 3; ≤days = 1), immunogenicity (low/privileged = 5; moderate = 3; very high = 1), and chemical resilience (very low sensitivity = 5; moderate = 3; very high = 1)^122–124^.

### 5.1 High-PPS Tier (15–18)

Ideal for durable bioengineering: **HIP-iPSCs (B2M**◼**/CIITA**◼**/CD47**◼**; 18)**—first clinical proof achieved with Sana Biotechnology’s UP421 in T1D patient, C-peptide production at 12 months without immunosuppression^228^; HSC HLA-matched (16; 5,000–30,000 clones contributing decades post-transplant^246^); autologous iPSC (16); chondrocytes (16; MACI grafts durable at 10–17 years^247^); astrocytes (16); Sertoli cells (16); engineered Tregs (15); microglia (15); NSC (15); neurons (15)^125–127^.

### 5.2 Upper-Mid Tier (12–14)

Kupffer macrophages (14), T-regs (14), memory B cells (14), MSC (14), γδ T cells (13; MHC-independent, no GvHD risk), naïve T cells (13), iPSC-NK (13), long-lived plasma cells (13), skeletal muscle fibers (13), hepatocytes (12), iNKT cells (12), CAR-T 4-1BB products (12; 96% detectable at 6 months vs. 65% for CD28 products^248^)^128–130^.

### 5.3 Mid Tier (10–11)

Alveolar macrophages (11), NK cells (11), cardiomyocytes (11), keratinocytes (10), pancreatic β-cells (10)^131–133^.

### 5.4 Low-PPS Tier (≤9)

Neutrophils (8), CAR-macrophages (9; limited by myeloid persistence), eosinophils/basophils (9), dendritic cells (7), effector T (7), endothelia (6–7), enterocytes (4)^134–139^.

**Table 1.** Mammalian Cell Type Characteristics Matrix (Updated 2025)

| Cell Type | Lifespan (in vivo) | Persistence (post-Tx) | Immuno-genicity | Chemical Sensitivity | PPS |
| --- | --- | --- | --- | --- | --- |
| Neutrophils | 19h–5.4d* | Hours | Low | Very High | 8 |
| Classical Monocytes | 1–3 days | Days | High | Moderate | 9 |
| Microglia | 4.2yr avg† | Years | Low–Mod | High | 15 |
| Dendritic Cells | 3–10 days | Days | Very High | High | 7 |
| NK Cells | ~2 weeks | Wk–months | High | Moderate | 11 |
| γδ T Cells‡ | Weeks–months | Weeks–mo | Low–Mod | Low–Mod | 13 |
| Naïve T Cells | 2–6 years | Autologous | Very High | Moderate | 13 |
| Effector T Cells | Days–weeks | Days–wk | Very High | Mod–High | 7 |
| Memory T Cells | Mo–years | Months | Very High | Low–Mod | 14 |
| Regulatory T Cells | Years | Months | Moderate | Moderate | 14 |
| Memory B Cells | Years | Months | High | Low | 14 |
| iNKT Cells‡ | Wk–months | Weeks | Low–Mod | Low–Mod | 12 |
| Hepatocytes (diploid) | <3yr avg† | Mo–years | Mod–High | High | 12 |
| Cardiomyocytes | Decades | Wk–months | Very High | Mod–High | 11 |
| Skeletal Muscle | 10–16 years | Mo–years | Very High | Moderate | 13 |
| Enterocytes | 3–5 days | Hours–days | Very High | High | 4 |
| Chondrocytes | Decades | Months | Moderate | Low–Mod | 16 |
| Neurons | Lifetime | Years | Low–Mod | High | 15 |
| Astrocytes | Years | Mo–years | Low–Mod | Moderate | 16 |
| HSCs (HLA-matched) | Decades | Years†† | Low | Very High | 16 |
| NSCs | Years | Months | Moderate | Very High | 15 |
| MSCs | Mo–years | Days–wk§ | Low | Moderate | 14 |
| iPSCs (autologous) | Indefinite | Variable | Moderate | High | 16 |
| HIP-iPSCs‡ | Indefinite | Years††† | Very Low | High | 18 |
| CAR-T (4-1BB)‡ | Engineered | >10 years§§ | High (allo) | Moderate | 12 |
| iPSC-NK‡ | Indefinite | Wk–months | Low–Mod | Moderate | 13 |
| Eng. Tregs‡ | Years | Months | Low–Mod | Moderate | 15 |
| CAR-Macrophages‡ | Days–weeks | Days–wk | Moderate | Moderate | 9 |

\*Lifespan debated: conventional 19h half-life vs. 5.4d total (deuterium labeling). †<sup>14</sup>C birth-dating data. ‡Newly added cell type. ††Based on 5,000–30,000 clones contributing 9–31yr post-HCT (Nature, 2024). §Paracrine “hit and run” mechanism; Ryoncil approved Dec 2024. †††Sana UP421: C-peptide at 12mo without immunosuppression (NEJM, 2025). §§Melenhorst et al.: CAR-T detectable >10yr post-infusion.

## 6. Advanced Profiling Technologies and Methodologies

Single-cell transcriptomics platforms like 10x Genomics Chromium enable analysis of thousands of individual cells simultaneously^140–142^. STAMP transforms imaging platforms into scalable profiling tools processing >10 million cells^143^. Organ-on-chip platforms enable species-specific toxicity assessment^148–153^. The new ISO/TC 276/SC 2 subcommittee on microphysiological systems is developing standardized cell sourcing criteria for organ-on-chip platforms^249^—a natural application for PPS-style scoring.

## 7. Clinical Applications and Regulatory Landscape

### 7.1 Regenerative Medicine and Cell Therapy

Clinical translation has accelerated significantly^154–156^. The Personalized Medicine Coalition’s 2025 report documents that ∼38% of FDA-approved drugs in 2024 required biomarker-guided administration—exceeding 25% for ten consecutive years^250^. The global cell analysis market was valued at approximately $32 billion in 2024, projected to reach $55.3 billion by 2030 at ∼10% CAGR^251^.

The FDA approved 8 new cell and gene therapy products in 2024, including AMTAGVI (first TIL therapy) and Ryoncil (first MSC therapy)^252^. As of March 2026, ∼47 total CGT products are FDA-approved. While the projected 10–20 annual CGT approvals by 2025 has not yet fully materialized (3–5 approvals in 2025), RMAT designations surged to 50 grants in FY2025—the highest ever—indicating growing regulatory appetite^253^.

### 7.2 Gene Editing and Cellular Engineering

Casgevy (exagamglogene autotemcel) was approved for SCD on December 8, 2023 and TDT on January 16, 2024, with 39 patients treated globally through Q3 2025 and approvals spanning the US, UK, EU, Canada, and Gulf states^161,254^. No additional CRISPR-based therapies have been approved since, though ∼250 gene-editing clinical trials are active worldwide. Beyond CRISPR nucleases, at least 15 base editing trials are active across 5 countries (BEAM-101 received RMAT designation), and the first-ever prime editing therapy (PM359, Prime Medicine) demonstrated safety and efficacy in a chronic granulomatous disease patient in May 2025^255^.

### 7.3 In Vivo Cell Programming

In vivo CAR-T generation represents perhaps the most disruptive advance for the PPS framework. Kelonia Therapeutics’ KLN-1010—the first-in-human in vivo CAR-T—achieved MRD-negative responses in all 4 treated multiple myeloma patients at month 1, with memory-phenotype CAR-T cells detected through month 3, requiring no lymphodepletion, apheresis, or ex vivo manufacturing^256^. A 2026 *Nature* publication demonstrated CRISPR-Cas9 site-specific CAR transgene integration in vivo^257^. These advances suggest the PPS should incorporate an “in vivo programmability” dimension reflecting whether cell types can be engineered directly within the body.

### 7.4 Digital Twins and AI Virtual Cells

The Chan Zuckerberg Initiative published a community roadmap for AI Virtual Cells in *Cell* (December 2024) with 40+ co-authors^173^. The CZI Virtual Cells Platform now hosts curated models including scGPT, scGenePT, and TranscriptFormer. CZI’s rBio reasoning model (2025), trained on virtual cell simulations, enables natural-language biological queries^258^. CZI partnered with NVIDIA in October 2025 to scale to petabytes of cellular data. For PPS, foundation models could eventually predict sub-scores from transcriptomic signatures—though a 2025 *Genome Biology* study cautioned that in zero-shot settings, scGPT may be outperformed by simpler methods for basic tasks^259^.

## 8. Regulatory Framework and Standardization

The FDA released directly relevant guidance: “Considerations for CAR T Cell Products” (final, January 2024), “Potency Assurance for CGT Products” introducing the potency assurance strategy concept, and “Safety Testing of Human Allogeneic Cells” (draft, April 2024)^260^. The EMA’s comprehensive ATMP guideline became effective July 1, 2025. The ICH Cell and Gene Therapy Discussion Group identified >300 ATMP-specific guidelines across member jurisdictions in December 2024, underscoring the fragmentation that a standardized scoring framework could help address^261^. The EU JRC published a roadmap for organ-on-chip standardization in January 2025^249^.

## 9. Current Limitations and Future Directions

### 9.1 Technical Challenges

Pipeline optimization remains complex^186–188^. 3D analysis capabilities lag behind 2D approaches^189^. Multimodal integration continues to limit temporal dynamics capture^190–195^.

### 9.2 New Lineage Tracing Techniques for Persistence Measurement

EPI-Clone (*Nature*, 2025) enables transgene-free clonal tracing using stochastic DNA methylation epimutations as digital barcodes, applied to 230,358 single cells of human hematopoiesis^262^. CRISPR barcoding advances include SMALT (deaminase-based substitution mutations), DAISY-barcode + Cas12a tracking >47,000 melanoma cells, and CellTag-multi for single-cell lineage capture across modalities (*Nature Biotechnology*, 2024)^263^. These tools make the persistence dimension of PPS more directly measurable than when the framework was conceived.

### 9.3 Research Priorities

Critical needs include: development of standardized PPS benchmarks validated against clinical outcomes; integration of in vivo programmability metrics; real-time non-destructive monitoring technologies; and computational tools that leverage foundation models for automated sub-score prediction from transcriptomic signatures^205–210^.

## 10. Discussion

This profiling reveals fundamental trade-offs best visualized through Pareto frontier analysis (Figure 5): short-lived innate cells excel in acute detection but falter in persistence, while privileged CNS/testis cells enable long-term grafts but resist engineering^211–216^. The Pareto framework reveals that HIP-iPSCs approach the theoretical optimum—combining indefinite expandability with immune evasion proven in humans—while γδ T cells and iPSC-NK cells occupy previously uncharted positions combining allogeneic compatibility with moderate persistence.

The linear PPS remains a practical first-pass screening tool, but we recommend supplementing it with: (a) Pareto frontier visualization for multi-objective analysis, (b) fuzzy membership functions for parameters with high measurement uncertainty (particularly neutrophil lifespan and hepatocyte heterogeneity), and (c) sensitivity analysis across weight configurations. The emergence of in vivo CAR-T generation^256,257^ further suggests a fifth dimension—in vivo programmability—that could fundamentally alter scoring for accessible immune populations.

## 11. Conclusions

The PPS scoring system occupies genuine white space—no competing composite index exists for cross-cell-type comparison—and offers a practical tool for researchers and clinicians. By extending the framework with Pareto analysis, emerging engineered populations (HIP-iPSCs, γδ T cells, in vivo CAR-T), and mathematical alternatives to linear scoring, we transform cell-type selection from empirical judgment into a reproducible, design-oriented process. As foundation models mature, PPS sub-scores may eventually be predicted from transcriptomic profiles, enabling real-time cell selection optimization for precision medicine.

## Acknowledgments

This synthesis is based on compiled datasets from cell profiling, scoring, and stem cell documents. The authors acknowledge the broader scientific community in advancing cell characterization methodologies.

## Declaration of Generative AI and AI-Assisted Technologies

During the preparation of this work the author(s) used ChatGPT in order to find citations as well as reorganize topics in the manuscript after drafting for logical flow. After using this tool/service, the author(s) reviewed and edited the content as needed and take(s) full responsibility for the content.

## Declaration of Interests

All authors are employed by Ainnocence Inc. The corresponding author is the founder of Ainnocence Inc. There are no patents or external funding sources to declare.

## Notes

### Competing Interest Statement

The authors have declared no competing interest.

